# Pattern separation is the key driver of expectation-modulated memory

**DOI:** 10.1101/577791

**Authors:** Darya Frank, Daniela Montaldi

## Abstract

The hippocampus uses pattern separation and pattern completion in a continuous manner to successfully encode and retrieve memories^1,2^. However, whether and how cognitive factors might modulate the dynamics between these types of computation is not well understood. Here we examine the role of expectation in shifting the hippocampus to perform pattern separation. Expectation can be built up through multiple contextual exposures leading to prediction (as in a learnt contingency) or through logical deduction based on a previous mnemonic response. Participants first learned a contingency between a cue and an object’s category (man-made or natural). Then, at encoding, one third of the cues that preceded the to-be-memorised objects violated the studied rule. At test, participants performed an old/new recognition task with old items (targets) and a set of parametrically manipulated (very similar to dissimilar) new foils for each object. We explored the effects of both contextual expectation, manipulated at initial encoding, and mnemonic-attribution expectation, driven by the mnemonic decisions taken on previous retrieval trials. For example, a target would be unexpected if in a previous trial a similar foil had been erroneously accepted as old. Memory was found to be better for foils of high and mid similarity to contextually unexpected targets at encoding, compared to expected ones. Additionally, violations of mnemonic-attribution expectation also yielded improved memory performance when the level of foil similarity was high. These results suggest that violations of both contextual expectation and mnemonic-attribution expectation engage pattern separation, resulting in better discrimination performance for these items. Importantly, this mechanism is engaged when input differentiation is required in order to make a correct recognition decision.

## Introduction

Memory is a dynamic process; the information we encounter, as well as our internal state, can affect how we remember. This intrinsic characteristic makes episodic memory a key aspect of adaptive behaviour. Whether a given event or item will be successfully encoded and subsequently retrieved from memory will depend on multiple contributing factors, ranging from the outputs of lower-level biological computations to the products of complex higher-level cognitive processes. Memory experiences can change dynamically, depending on what we expect to encounter in everyday life and on its mnemonic attribution. For example, an unexpected sequence of events might make your commute to work today stand out from previous ones. Critically, what we expect to encounter may be driven by external cues (e.g. the contextual setting) but it can also be driven by internally generated cues (i.e. spontaneous recall of a previous memory experience). Whilst there is evidence for the effect of the former on memory performance, the latter has received little attention. Two questions therefore arise; what underlying memory mechanism renders unexpected encounters more recallable? And, can this mechanism be triggered by both externally- and internally-cued expectation? To answer these questions, we employ a novel approach to elicit both an externally-driven explicit rule-based expectation and an internally-driven dynamic mnemonic-attribution expectation. We examine whether they both engage a fundamental computation underlying memory; that of pattern separation.

As we constantly encode and retrieve information from memory, it is easy to overlook the potential complexity of the mechanisms supporting these processes. Perhaps the most computationally-demanding task is to separate memories of similar events. Consider trying to remember whether you have locked your door today. You probably have multiple experiences of locking your door, and therefore would need a distinct way to represent today as opposed to any other day. To achieve this, the hippocampus performs constant comparisons between existing and new information, resulting in pattern separation (PS); encoding the current input as separate and distinct from existing memories, and pattern completion (PC); retrieving a complete memory representation from a partial cue^1,3^. These computations are complementary^4^ and believed to occur incessantly^5–7^ to support episodic memory. If you have successfully pattern-separated locking your door today, it should be easy to retrieve this memory accurately. However, similar memories often interfere, leading to erroneous pattern completion to another stored memory. Pattern separating an episode is thus essential to being able to later successfully pattern complete (recall) it. Therefore, identifying external or internal cognitive factors that facilitate a shift towards PS could improve our understanding of what makes some memories more readily recallable than others.

Whilst dynamic shifts between PS and PC have been discussed extensively^2,3,5,7^, the possibility of top-down cognitive processes tilting the scale towards PS or PC has not been well studied. Such a modulator would need to compute discrepancies between current input and stored information, to capture mismatches or update the predictive value of existing representations^8^. One cognitive process that performs such comparisons is expectation. Indeed, top-down expectation plays a significant role in guiding adaptive behaviour^9–12^. Recently, there has been growing interest in how expectation drives memory processes^13–15^. Consequently, the advantageous effect of contextual surprise on memory has been observed in numerous studies^16–18^. The hippocampus, perhaps through interaction with the dopaminergic system^19–22^, is believed to support this effect by engaging an encoding state, leading to improved memory performance^21,23–25^. Findings from animal models provide some support for this claim, showing context-sensitive shifts between PS and PC^26,27^. However, evidence to date has failed to demonstrate the relationship between contextual expectation and pattern separation/completion markers in humans. To shed light on this relationship, comparison is needed between expected and unexpected items in tasks where memory performance is dependent on successful pattern separation.

Given that PS entails disambiguating similar inputs, whereas PC results in integrating across inputs, a common approach to probe them is using perceptually similar foils^28–31^. In such tasks, ‘old’ responses to foils (false alarms) can be interpreted as a result of erroneously pattern completing to target, whereas ‘new’ responses to foils (correct rejections) capture pattern separation, distinct from the target. As similarity increases between target and foils, discrimination becomes more difficult, therefore requiring pattern separation. If a violation of expectation triggers a shift towards a PS encoding state, items from sets whose target was unexpected would be more easily discriminated than those from expected sets, of the same similarity level. Alternatively, if expectation does not modulate memory, performance should reflect solely the degree of similarity. Using a discrimination task also allows us to directly test whether the beneficial effect of surprise *selectively* targets a pattern separation mechanism or provides a more general memory boost. If the latter is true, we should observe more hits as well as more correct rejections of all unexpected items (a global enhancement effect). On the other hand, an effect that is selective to correct rejections of unexpected similar foils would suggest violations of expectation trigger a shift towards PS. This mechanism would only be observed when input differentiation is essential to task performance.

Despite the fact that expectation can be elicited and violated in different ways, to the best of our knowledge, research to date has only focused on external (contextual) expectation^16–18^ and has not distinguished between different sources of expectation. However, employing a behavioural pattern separation task, with several foils of different levels of similarity to each target, also enables us to explore mnemonic-attribution expectation. This expectation, which we believe to be predominantly implicit, is driven by the mnemonic status attributed to existing memorial representations. Unlike explicit expectation, which can be prompted using a learned contingency^12^ and requires an external contextual cue, mnemonic-attribution expectation is elicited internally and therefore may be less likely to require conscious processing. In everyday life we do not normally explicitly label information as old or new, as we would in a lab-based task. Nevertheless, given the dynamic nature of memory, these processes still take place, implicitly, as one stimulus representation can evoke a memory representation of a previous stimulus. Such ‘automatic memory recording’^32^ has been used to explain binding of items to context, which forms the basis for the contextual expectation, introduced above. Here, we extend this notion to include not only the item itself, but also its attributed mnemonic status. Therefore, mnemonic-attribution expectation would encompass the conjunction between a sequence of items at retrieval, and their mnemonic representation (whether they were previously judged as old or new).

In the current study, therefore, we refer to mnemonic-attribution expectation as that arising from previous mnemonic responses to similar items presented within the retrieval phase. For example, a foil prompting an ‘old’ response (false alarm) would make a subsequent target *unexpected*, as the target had already been identified (albeit erroneously). Mnemonic-attribution expectation requires reinstatement of the decision made in an earlier trial, which can then be used to guide the current memory judgment^31^. It is therefore more dynamic, and dependent on the continuous retrieval of previous mnemonic decisions, rather than on a single pre-defined context. It thus follows that the choice of retrieval task in this case is crucial; a continuous old/new (yes/no) recognition task is best suited to examine such effects as it allows independent mnemonic decision for each stimulus^33^ (i.e. no restriction on ‘old’ responses for stimuli from the same set). If a previous mnemonic decision has elicited an expectation (which can be met or violated during the current trial), we should observe differences in memory performance depending on whether correct or incorrect previous responses were made. By combining mnemonic-attribution expectations and contextual expectations in one study, we can explore, more fully, the extent to which different sources of expectation modulate memory, while shedding light on the nature of any underlying mechanisms.

Here we developed a novel behavioural approach that allows us to test whether the beneficial influence of expectation on memory is driven by a shift towards PS, and to explore the specificity of this memory-enhancement mechanism. To elicit contextual expectation, we employed a rule-learning task where participants learned an association between a cue and a category (man-made or natural). In each trial, participants were presented with a cue for 1s and asked to guess the following item’s category. They then received feedback about their decision and were instructed to use it to learn the contingency between the cue and category. In two encoding runs, participants were presented with the same set of cues, followed by an object to encode. In the first round all cues were rule-abiding, however, in the second round, some of the cues were incongruent with the subsequent stimulus, resulting in *unexpected* encoding trials. At retrieval, we employed foils of parametrically-manipulated similarity (high similarity F1, mid similarity F2, low similarity F3, using stimuli and dissimilarity indices provided by SOLID^34^). Targets and their three associated foils formed sets; the old/new recognition task allowed the modulation of mnemonic-attribution expectation, elicited by decisions made on previous presentations of items from the same set. This paradigm (see Figure 1A-C) allowed us to address two questions; first, does expectation affect memory performance by modulating shifts between pattern separation and pattern completion? Secondly, are expectation effects specific to contextual expectations, or can they also be driven by a more dynamic phenomenon involving the comparison between previous mnemonic decisions (experiences) and current inputs (i.e., mnemonic-attribution context)? We hypothesised that similar discrimination performance would be observed for both contextual and mnemonic-attribution expectations. Specifically, we predicted that unexpected items would benefit from enhanced discrimination performance compared to expected ones, driven by a shift towards PS.

**Figure 1.**
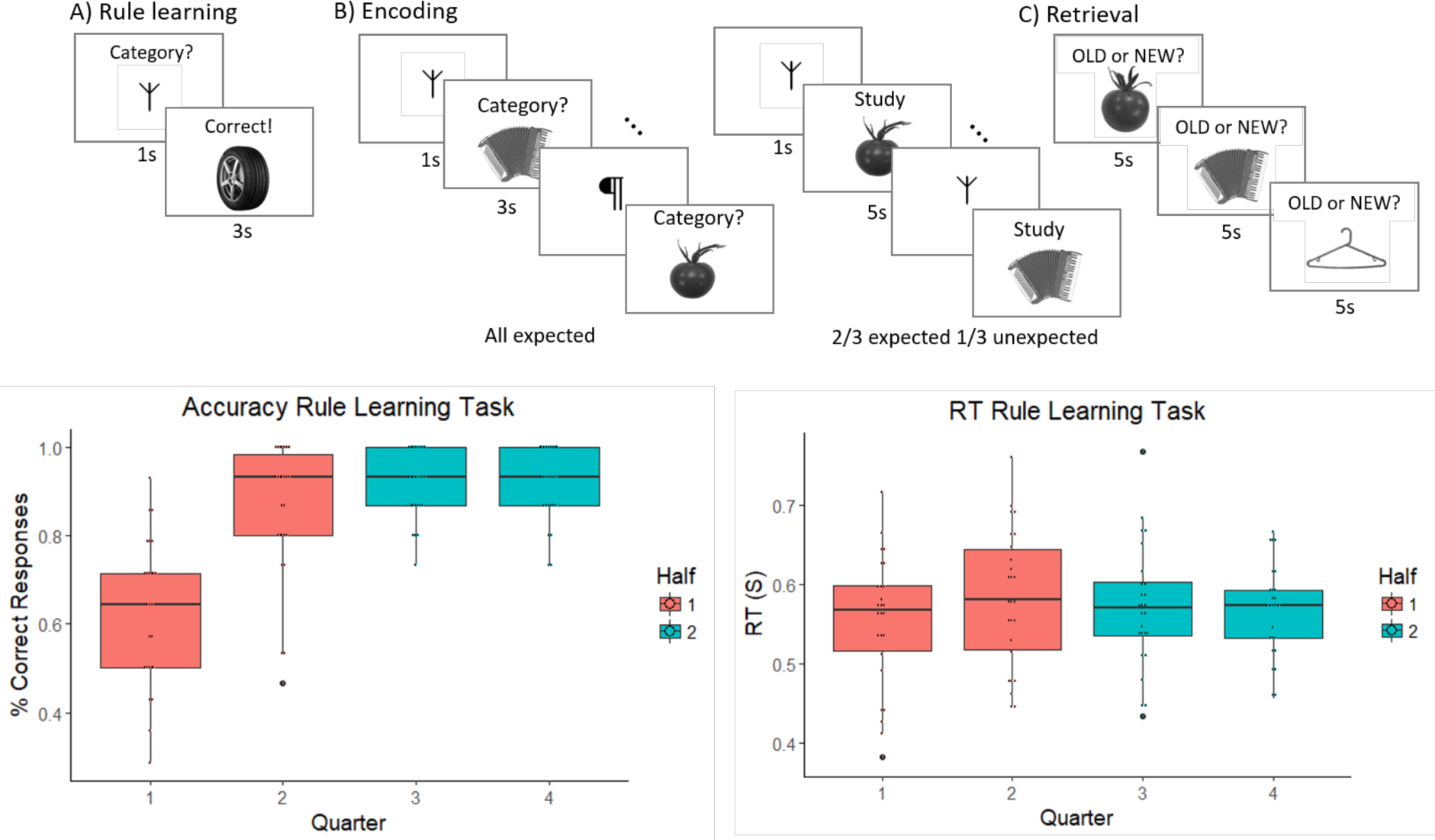
Experimental design and rule learning task results. A) In the rule-learning task participants learned a contingency between a cue and an object’s category, man-made or natural. B) During the first round of encoding, participants were presented with the same cues (all rule-abiding) and had to indicate whether the object was man-made or natural. In the second round, participants are asked to study the item carefully and 1/3 of the cues are misleading (unexpected). C) In the final retrieval task old items (targets) and new similar foils are presented, and participants are asked to respond whether the item is old or new. D) Accuracy in rule learning task as a factor of task progression. Participants learned the cue-category contingency well and their accuracy was above chance from the second half of the task. E) Reaction time (RT) was restricted to 1s, so no differences in mean RT were observed, but there was a reduction in variance as the task progressed and participants learned the contingency. Unless otherwise stated, error bars in all figures reflect standard error; * p ≤ 0.05, ** p ≤ 0.01, *** p ≤ 0.001.

## Results

### Rule-learning performance

To ensure contextual expectations were established, we examined participants’ memory performance following the second half of the rule-learning task (Figure 1D-E). Average performance, excluding two participants who did not reach criterion (75% accuracy), was 91.6% (SD = 6.5%), significantly above chance (t(25) = 32.77, p < 0.001, Cohen’s d = 6.42). Next, we examined reaction times for the predictions, within the 1s decision window. There was no significant difference in mean reaction time between the first and second halves of the task (t(25) = 0.001, p = 0.999).

### Recognition memory performance

To quantify the probability of giving a correct response on each retrieval trial we used mixed-effects logistic regression modelling. This allowed us to capture the influence on memory, of the contextual expectation set at encoding, as well as the expectation driven by previous mnemonic decisions for objects from the same set, and the distance (number of intervening trials) between previous and current trials.

As contextual expectations were manipulated at encoding, we can look at the first retrieval presentation of an item to examine contextual expectation effects without any interference driven by mnemonic-attribution expectation (Figure 2). There were more correct rejections of high similarity foils (F1s) of unexpected targets, compared to expected ones (β = 0.466, z = 2.4, p = 0.016). Mid-similarity foils (F2s) showed a similar trend, with more correct rejections for unexpected F2 from sets whose target was unexpected at encoding (β = 0.405, z = 1.92, p = 0.054). Targets (β = 0.181, SE = 0.339, z = 0.523, p = 0.592) and low similarity foils (F3s) (β = −0.118, z = −0.533, p = 0.594) did not show any contextual expectation effects. These results therefore show a selective increase in the correct rejection of foils similar to contextually unexpected targets.

**Figure 2.**
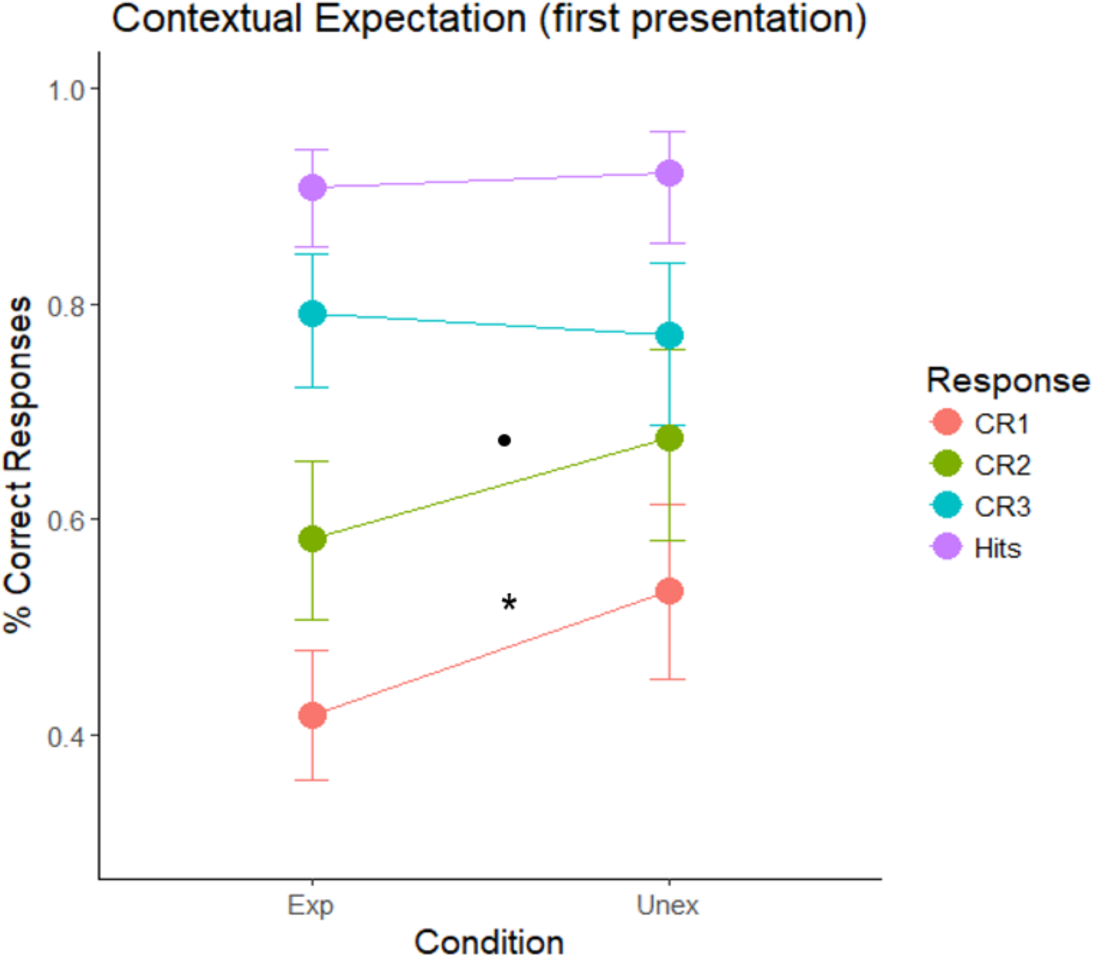
Contextual expectation. Contextually unexpected high (F1) and mid (F2; p = 0.054) similarity foils were correctly rejected (CR1, CR2) more than expected ones. No differences were observed for low similarity foils (F3) or targets.

Next, we sought to explore how mnemonic-attribution expectation affected memory performance and whether it interacted with contextual expectation. To achieve this, we extracted items that were presented later in the set position at retrieval, after all other items from the same set had already been presented and attributed a mnemonic decision. For clarity, we report here the results for the last items (4^th^ position in the set), similar results were obtained using unique models for each of the preceding items (see Supplementary Materials). When targets were presented last in the set sequence (Figure 3), we observed more hits following a false alarm response to F1 (β = −1.44, z = - 3.38, p < 0.001) and F3 items (β = −0.752, z = −2.27, p = 0.023), compared to correct rejections. This shows that erroneous mnemonic attribution of previous F1 and F3 items as old was associated with more hits for following targets. No difference between F2 responses was observed (β = −0.272, z = - 1.14, p = 0.255). There was also a significant interaction between F1 response and distance between target and F1 (β = 2.04, z = 2.28, p = 0.022), showing there were more hits for targets following false alarms of high similarity foils (FA1), compared to CR1, when targets and F1 were closer in the retrieval phase.

**Figure 3.**
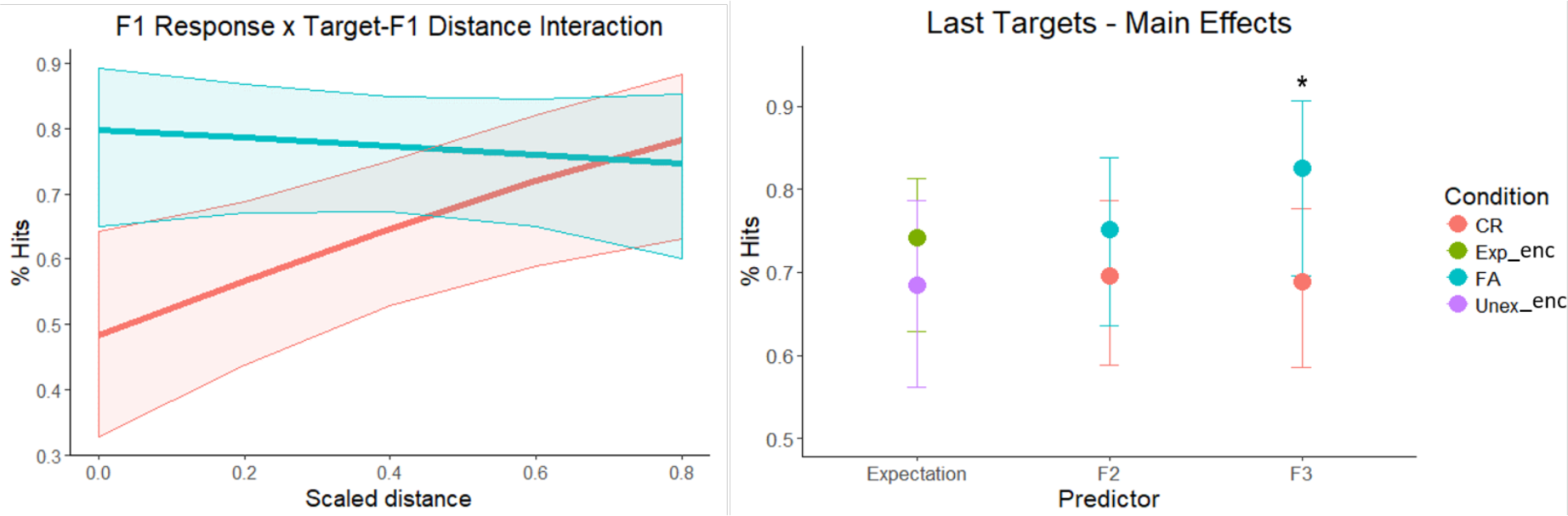
Predicting hits. A) There were more hits for targets following false alarms of high similarity foils (FA1), especially if they were closer in time. B) More target hits were observed following low similarity false alarms (FA3). Target’s contextual expectation (set during encoding) and F2 response did not affect hits.

Next, we examined F1 trials that were presented last to evaluate the expectation effect, elicited by mnemonic attributions of previous items from the same set, at retrieval (Figure 4a). We observed more correct rejections following missed targets than following hits (β = −1.02, z = −3.79, p < 0.001), indicating a mnemonic-attribution expectation effect of previously missed targets on correct rejections of current F1 items, rendering them unexpected. Higher levels of CR1 were also observed following previous correct rejections of F2 (β = 0.814, z = 3.89, p < 0.001) and F3 (β = 0.774, z = 3.15, p = 0.002), compared to false alarms. Including distances between F1 and any of the previous items from the same set did not significantly improve the model (χ^2^ (3) = 0.255, p = 0.968). Correct rejections of last F2 items (Figure 4b) were boosted when they were preceded by correct rejections of F1 (β = 0.544, z = 2.3, p = 0.021) and F3 (β = 0.799, z = 2.86, p = 0.004), compared to false alarms. The effects of contextual expectation (β = 0.103, z = 0.429, p = 0.668) and target response (β = - 0.199, z = −0.648, p = 0.517) were not significant. Including distances between F2 and the previous items did not significantly improve the model (χ^2^ (3) = 2.11, p = 0.549). Finally, when F3 was presented last (Figure 4c), we observed more correct rejections when it was preceded by correct rejections of F2 compared to false alarms (β = 1.21, z = 3.49, p < 0.001). There were no significant effects of contextual expectation (β = −0.268, z = −0.774, p = 0.439), target (β = 0.122, z = 0.274, p = 0.784), or F1 responses (β = −0.02, z = −0.057, p = 0.954). Including distances in retrieval trials from previous items did not improve model fit (χ^2^ (3) = 5.6 p = 0.132).

**Figure 4.**
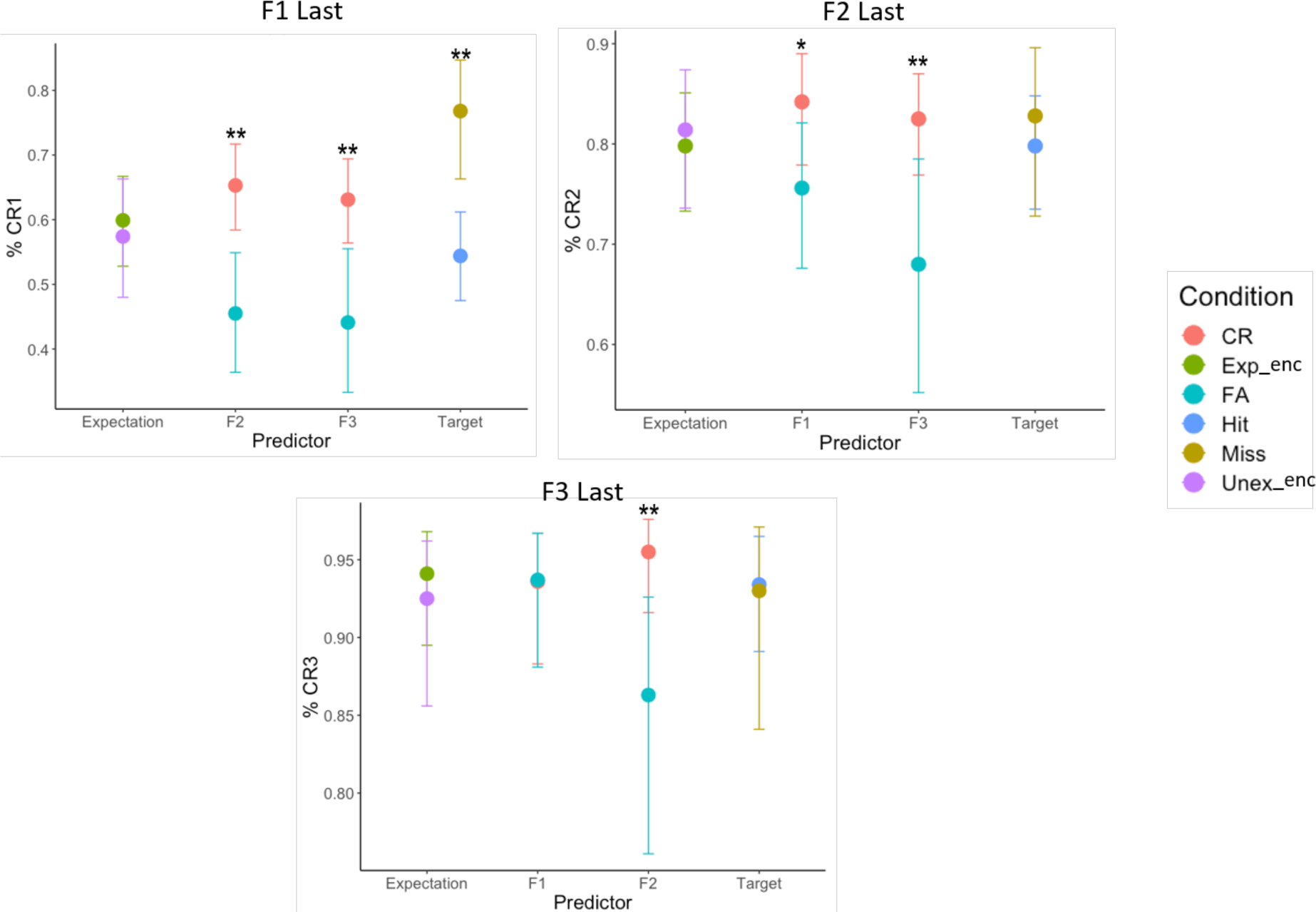
Predicting correct rejections. A) More correct rejections of high similarity foils (CR1) were observed when they were preceded by CR of other foils (CR2, CR3), and by missed targets. B) More correct rejections of mid similarity foils (CR2) were observed when they followed correct rejections of other foils (CR1, CR3). C) Low similarity foils had more correct rejections (CR3) when they were preceded by CR2.

## Discussion

This set of results offers the first direct evidence for the role of expectation in engaging a pattern separation mechanism. We found the contextual manipulation exerted an effect only on the first item presented within a set, when there was no mnemonic-attribution expectation. High and mid-similarity foils from sets whose target was unexpected at encoding produced more correct rejections at retrieval, compared to foils whose target was expected at encoding. Targets, as well as low similarity foils, were unaffected by this manipulation. As the retrieval task progressed, the effect of contextual expectation diminished and was replaced by the more-immediate mnemonic-attribution expectation. By examining the last items presented within a set at retrieval, we found hits were more likely following FA1 and FA3. A complementary effect was observed for CR1, with more correct rejections following misses, compared to hits. Finally, across similarity levels, the level of correct rejections of previous foils was associated with better discrimination performance (correct rejection of current foil). Overall, our results suggest expectations, both contextual and mnemonic-attribution, engage the hippocampal system and support memory performance by employing PS to disambiguate highly similar items. Such a mechanism could support adaptive memory, making unexpected information more memorable.

An abundance of literature has addressed hippocampal shifts between PS and PC, mostly in rodents^26,27^, and as a possible factor contributing to deteriorating memory in ageing^35,36^. However, the contribution (if any) of top-down cognitive factors to these shifts has remained unclear. If the balance between PS and PC was controlled merely by bottom-up information, based on a raw input similarity function, we would not expect a cognitive manipulation to make any difference. Instead, here we show top-down expectations, both contextual and mnemonic-attribution expectations, play a role in engaging PS. Unexpected similar items were more accurately recognised than expected ones, whereas there was no expectation-driven difference in memory performance for items of a lower similarity. It is important to note that although we did not record neural activity in this study, our experimental design explicitly taps behavioural markers of pattern separation^28–31^, which supports recall and is heavily dependent on hippocampal function^4^.

Previous studies have shown that contextual expectation plays an important role in modulating hippocampal involvement and behavioural memory responses^13,18,22,23^. However, the extent of these effects, and the underlying mechanism supporting them, remained unclear. Here we show contextual surprise does not enhance memory unequivocally, but specifically aids the disambiguation of overlapping inputs. This suggests that the beneficial effect of surprise stems from enhanced PS rather than a more general memory boost, which could be mediated by an extra-hippocampal circuit. We found that violations of expectation at encoding support the ability to correctly identify similar foils as such. Additionally, we did not observe any beneficial effect of contextual expectations on hits or correct rejections of low-similarity foils. Taken together, these results strongly suggest that a modulating mechanism is employed when unexpected events occur, leading to a shift towards pattern separation of the surprising information. Importantly, the behavioural output of such a mechanism depends on the amount of overlap between existing and new inputs. In cases of full input-test overlap (targets) and low overlap (low similarity foils), pattern separation is redundant in supporting memory performance, therefore even if the shift occurs, it would not be reflected in the memory response. Our results are consistent with previous computational work^4^ showing hippocampal PS peaks at high (but not full) input similarity level, and gradually decreases as similarity levels advance towards either end of the scale (see Figure 5).

**Figure 5.**
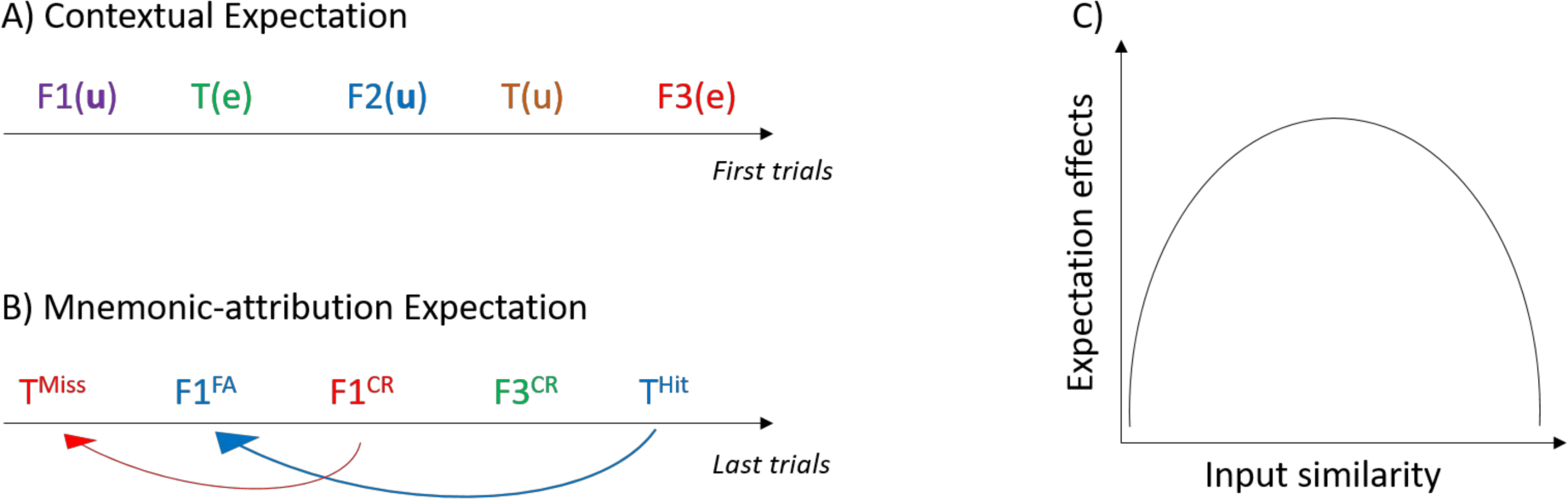
Conceptual Illustration. A) Contextual expectation generated at encoding. Each encoding trial was associated with a cue from the rule learning task. An item is unexpected (u) when there is a mismatch between cue and the item’s category and expected (e) when there is no mismatch. At retrieval, the independent effect of this expectation can be examined in the first presentation of items from unique sets. Different sets are represented by different colours. B) Mnemonic-attribution expectation generated at retrieval. The response given to each item is in superscript, arrows reflect the mnemonic-attribution expectation within a set. When a previous response was incorrect (e.g. miss) the following presentation of a similar item from the same set prompts violation of the mnemonic-attribution expectation and a subsequent correct response (e.g. CR). C) Illustration of the memory effects. When input similarity between encoding and retrieval is very low (target-F3) or very high (target-target), violation of expectation does not exert an effect on memory. When similarity is sufficiently high (e.g. target-F1), violation of both contextual and mnemonic-attribution expectations boosts memory performance.

A PS-dependent effect was also observed for mnemonic-attribution expectation. This expectation was elicited dynamically during retrieval and changed according to participants’ mnemonic decision on previous trials. Here, if a previous foil elicited an ‘old’ response (erroneously identified as a target), the subsequent presentation of the target from the same set would be unexpected. Indeed, we found in trials like this, unexpected items were associated with better memory performance. Importantly, this was only true for targets and high similarity foils, whereas mid and low similarity foils were unaffected. These results suggest that the continuous comparison between current and previous inputs elicits a similar effect to that of contextual expectation, supported by increased PS of unexpected items. While the notion that memory is altered by information presented at retrieval, is not a new one^37^, it has been discussed predominantly in relation to the presentation of semantic features or external associative cues as modulating factors^38^. Here, we show that the mnemonic attribution of previous items (hit, miss, CR or FA) also plays an important role in modifying memory performance. The mnemonic-attribution expectation demonstrated here reflects the continuous re-encoding of test items based on the mnemonic decision their set members previously elicited, combined with their perceptual similarity. This finding pertains to the dynamic nature of expectation and its resulting influence on memory decisions.

Despite the fact that our recognition task encouraged an independent mnemonic decision on each retrieval trial, we found that previous decisions modulated current ones, dependent on the level of perceptual similarity between the items. We interpret this finding as a demonstration of an implicit form of expectation that tracks not only the dynamic sequence of stimuli presented, but also their mnemonic attribution. An alternative account could be related to online error awareness; for instance, if a target is missed and the participant realised their recognition decision was wrong in real time, we would predict all following foils to be correctly rejected. However, our results show a specific effect of subsequent correct rejections of F1, but not F2 and F3 (see Supplementary Figure 1 as well). This pattern also negates the possibility that memory performance for items from the same set could be driven by enhanced encoding of the target, in which case we would predict equivalent response accuracy across items. Instead, these findings provide novel evidence for the important contribution that implicit mnemonic processes play in other areas of cognition, such as decision-making^39^. They could help elucidate how previous experience, and its memorial representation (including its mnemonic attribution), implicitly affects external markers such as explicit choices. To this end, future research would also benefit from examining how mnemonic-attribution expectation is manifested over a longer time scale (e.g. over days).

Although our results indicate that contextual and mnemonic-attribution expectations show a similar effect, possibly supported by a common computation of expected vs. received input^8,18,19^, we did not observe an interaction between the two types of expectation. Given that contextual expectation was manipulated only at encoding, whereas mnemonic-attribution expectation dynamically changes at retrieval, this is not surprising. Nevertheless, the lack of interaction does offer insight into the sensitivity of expectation effects on memory performance. In the current study, mnemonic-attribution expectation was object-specific, whereas contextual expectations were binary, depending on the item’s category (man-made or natural). A recent study^25^ showed increased hippocampal mismatch signals with increased prediction strength, and prediction-outcome similarity. Therefore, the stronger effect exerted by mnemonic-attribution expectation in our study could be explained by the higher degree of similarity between outcome and prediction, with object-dependent predictions being more specific (and thus more similar) than category-dependent ones.

In conclusion, we provide the first evidence for the modulating role of expectations on the behavioural marker of hippocampal pattern separation. We introduce the concept of mnemonic-attribution expectation, and show that violations of both this form of expectation and contextual expectation result in enhanced memory discrimination performance, supported by pattern separation. The specificity of this effect to highly overlapping inputs provides further support for the role of PS as the driving force behind the enhancement effects of surprise on memory. Future research should examine interactions between contextual and mnemonic-attribution expectations during retrieval, and their neural correlates.

## Methods

### Participants

29 participants (mean age = 19.5, 7 males) gave informed consent and took part in the experiment. Three participants were excluded from analysis due to memory performance below chance (one participant) or failure to perform the rule learning task (two participants). All procedures were approved by the University of Manchester Research Ethics Committee.

### Materials

78 images, natural (39) and man-made (39), were selected using the Dissimilarity Index from the Similar Object-Lures Image Database^34^ (SOLID). These images were used as the target objects, presented during encoding. Using a custom MATLAB (MathWorks Inc.) function (https://github.com/frdarya/SOLID/blob/master/ChooseFoils.m), three foils of decreasing levels of similarity were selected (DIs 1300, 2000 and 2700) for each target image. Similarity was parametrically manipulated by keeping the average distance between the levels constant (average DI between foils 2100). For the rule learning task 56 more images (28 from each manmade/natural category) were taken from SOLID.

### Procedure

The experiment was controlled using PsychoPy2 version 1.82^40^ and consisted of three main parts (see Figure 1a-c), similar to the design used by Kafkas and Montaldi^17^. **A) Rule learning task**. Participants learned an association between a symbol cue and a category (man-made or natural), to generate contextual expectations. There was a total of four cues, two for man-made and two for natural objects. Following a fixation cross, a cue appeared on the screen for 1s, during which participants were asked to predict the category of the next item. They were instructed to guess during the first few trials, but to learn the contingency as the task progressed. Subsequently, a man-made or natural object (not tested) appeared and participants received feedback about their prediction (3s). Each cue was repeated 14 times and cues were counterbalanced across participants.

**B) Encoding task.** In this task participants were presented with the 78 target images, encoded twice, in two rounds. During the first round, a previously-learned cue (1s) was followed by an object (3s), and participants were asked to indicate whether the object is man-made and natural. In this round all cues were consistent with the rule (no expectation violation). In the second round, participants were asked to study the perceptual details of the image carefully (5s). Importantly, 70% of the cues in this round were consistent with the rule (expected stimuli) while the other 30% of cues violated the rule (unexpected stimuli). In both encoding rounds participants were instructed to ignore the cue and focus on the main task. Stimulus presentation order was random, and allocation to expectation condition pseudo-random, maintaining equal number of expected-/unexpected items for the two categories. Prior to the retrieval task, an arithmetic distractor task was used for 5 minutes. **C) Retrieval task.** The final task was a continuous recognition memory paradigm. A stimulus appeared on the screen for 5 seconds during which participants had to decide if it was old (target) or new (foil).

### Data Analysis

Given the nature of our recognition task, and our interest in interactions between contextual and mnemonic-attribution expectations, we collated object sets (target + 3 foils) and ran mixed effect binary logistic regression models on these ungrouped data. Models were computed using the lme4 package^41^ in the R environment^42^. The parameters of such models can be used to assess the probability of giving a correct response (‘old’ for targets, ‘new’ for foils) whilst accounting for each participant’s unique intercept.

To assess the independent effect of contextual expectations established during encoding, we used a simple model using expectation as the only predictor, for each item (target, F1 etc.) separately. To examine the effect of mnemonic-attribution expectation we had to take into account the response to a previous item (hit, miss, FA or CR), and the distance (in number of trials) between these two items. We used a nested modelling approach, with predictors: contextual expectation, previous mnemonic response, and distance, and used a likelihood ratio test to identify the best performing model. This method examines the contribution of each a-priori chosen parameter to the overall model fit. Using the Akaike Information Criterion (AIC) to compare these models yielded identical results.

## Acknowledgments

The authors would like to thank Lewis Fry and Harry Hoyle for their assistance in data collection, and Alex Kafkas for helpful discussions. D.F. is supported by a PDS award from The University of Manchester.

## Authors Contribution

D.F. and D.M. designed the experiment and wrote the manuscript. D.F. collected and analysed the data.

